# Single-particle cryo-EM structures from iDPC-STEM at near-atomic resolution

**DOI:** 10.1101/2021.10.12.464113

**Authors:** Ivan Lazić, Maarten Wirix, Max Leo Leidl, Felix de Haas, Maximilian Beckers, Evgeniya V. Pechnikova, Knut Müller-Caspary, Ricardo Egoavil, Eric G.T. Bosch, Carsten Sachse

## Abstract

Electron cryo-microscopy (cryo-EM) is becoming one of the routine tools for structure determination of biological macromolecules. Commonly, molecular images are obtained by conventional transmission electron microcopy (CTEM) using underfocus and subsequently computationally combined into a high-resolution 3D structure. Here, we apply scanning transmission electron microscopy (STEM) using the integrated differential phase contrast mode also known as iDPC-STEM to the cryo-EM test specimen of tobacco mosaic virus (TMV). The micrographs show complete contrast transfer to high resolution and enable the cryo-EM structure determination at 3.5 Å resolution using single-particle based helical reconstruction. A series of cryo-EM TMV maps was resolved at near-atomic resolution taken at different convergence semi-angle (CSA) beams and share identical features with maps obtained by CTEM of a previously acquired same-sized TMV data set. The associated map B-factors from iDPC-STEM match those obtained by CTEM recordings using 2^nd^ generation direct electron detection devices. These data show that STEM imaging in general, and in particular the iDPC-STEM approach, can be applied to vitrified single-particle specimens to determine near-atomic resolution cryo-EM structures of biological macromolecules.

## Introduction

Scanning transmission electron microscopy (STEM) is a well-established methodology in characterizing materials at micro, nano and atomic scale [1, 2, 3]. A particular STEM method derivate known as ptychography was shown to image dose-resistant specimens reaching resolutions better than 0.5Å [4, 5] and is becoming the method of choice for the obtaining highest possible spatial resolutions [6]. Among these STEM imaging modalities, integrated differential phase contrast STEM (iDPC-STEM) [7, 8] has been routinely applied to a variety of specimens such as GaN, NdGaO_3_-La_0.67_Sr_0.33_MnO_3_, Ni-YSZ interfaces and Bi_2_Sr_2_CaCu_2_O_8+δ_ superconductors [9, 10, 11, 12]. In the same manner, specimens such as metal hydrides were successfully visualized at sub-atomic resolution, including heavy elements alongside light elements such as hydrogen [13]. Moreover, iDPC-STEM has been demonstrated to successfully image different crystalline as well as amorphous materials including beam-sensitive ones like zeolites [14, 15, 16]. One of these samples includes an individual aromatic hydrocarbon molecule trapped within a porous framework structure [17]. Other investigated materials are known as metal-organic frameworks (MOFs) that can only be imaged using electron doses smaller than 50 e^-^/Å^2^ before damaging the structure [18]. Using iDPC-STEM with a low-dose exposure of as little as 40 e^-^/Å^2^, a resolution of 1.8 Å has been successfully obtained from a single micrograph for such materials [19]. For these dose-sensitive low-contrast specimens, iDPC-STEM enables direct interpretation of the image without the need of defocusing and subsequent contrast-transfer function (CTF) correction of the images [20].

One of the early STEM applications to freeze-dried biological samples was the molecular mass determination by annular dark field (ADF) scattering, as the number of atoms is directly related to the scattering intensity [21]. More recently, STEM tomography has been applied to thick vitrified cells [22, 23]. Although low in resolution, it was shown that image contrast can be obtained even from μm-thick samples mainly by inelastically scattered electrons [24]. Furthermore, cryo-STEM has been applied to single-particle specimens of Fe or Zn-loaded ferritin to precisely visualize and locate metals within the protein cage [25]. More recently, micrographs of rotavirus and HIV-1 virus-like particles were imaged using ptychography reporting the principal application to biological specimens albeit at low resolution [26]. Thus far, complete 3D single-particle cryo-EM structures of biological macromolecules at close-to-atomic resolution, have not been determined by images of the STEM technique.

Electron cryo-microscopy (cryo-EM) has become a very successful structural biology technique that commonly produces near-atomic resolution 3D structures of ice-embedded biological macromolecules. Micrographs are obtained by conventional transmission electron microscopy (CTEM) at low fluences of 20 – 100 e^-^/Å^2^ and imaged at μm-defocus to boost low-resolution contrast of the macromolecules. When images of proteins assembled in regular 2D or helical arrays were available, they could be successfully determined at high resolution to enable atomic model building decades ago [27, 28, 29]. Due to the advent of improved hardware and software, also known as the resolution revolution [30], single-particle structures are now routinely resolved at near-atomic resolution. Thus, hundreds to thousands of micrographs are acquired at varying underfocus to increase the contrast of the ice-embedded macromolecules. Subsequently, elaborate image processing work-flows are employed to process the molecular projections, CTF correct and determine their orientation parameters to constructively merge often more than 10,000 of molecular views in a final 3D image reconstruction [31, 32]. At present, structures are commonly resolved better than 4.0 Å, suitable for atomic model building. To benchmark the performance of the cryo-EM method, the obtained resolutions of various test specimens such as tobacco mosaic virus (TMV) were improved over time to better than 2.0 Å [33]. So far, the highest resolutions of 1.2 Å were accomplished using another common test specimen known as apoferritin, [34, 35].

Due to the reported benefits of STEM approaches for a large range of different materials, we wanted to explore whether STEM imaging can be applied to vitrified biological samples and produce high-resolution images. Using iDPC-STEM imaging of TMV, we here demonstrate the successful single-particle based helical reconstruction at 3.5 Å resolution using an electron beam of 4.0 mrad convergence semi-angle (CSA). The resulting cryo-EM map matches the expected features of previously analyzed CTEM TMV data sets at the given resolution. The obtained quality of the iDPC-STEM map is comparable with those obtained with CTEM approaches recorded using 2^nd^ generation direct electron detection (DED) cameras [36]. This study shows that iDPC-STEM imaging can be successfully applied to cryo-EM single-particle based structure determination and elucidate biological structures at near-atomic resolution.

## Results

In order to successfully image vitrified samples by STEM, several additional aspects need to be considered in comparison with CTEM defocus-based imaging. First, instead of flood-beam illumination of a large field of view, the convergent beam moves over the specimen in regular intervals and illuminates the sample spot by spot to scan the region of interest (**Fig. 1a-b**). For each beam position of the scanning process, the resulting signal is recorded in the far field behind the sample using a center-of-mass (COM) detector, here approximated by means of a four-quadrant detector [7, 8]. Second, whereas in CTEM the optical resolution limit is caused by an inserted aperture and lens aberrations, in STEM the CSA of the focused beam controls the apparent resolution and the depth of focus (**Fig. 1c**) and aberrations, if present, will affect only the CTF shape but not limit the resolution [8]. Third, unlike CTEM imaging methods that produce additional phase contrast by defocusing the specimen, STEM techniques such as annular dark field (ADF) STEM and iDPC-STEM reach the highest contrast of the image in focus. Therefore, focus and maximized contrast was obtained by examining the flatness of the convergence beam diffraction pattern (CBED) of the beam (**Fig. 1d-e**). Under the described STEM imaging conditions (**Fig. 1f**), the typical field of view (FOV) of a cryo-micrograph is around 500 nm. In analogy to existing cryo-EM low-dose acquisition protocols, we avoid multiple exposures of the molecules and applied the focusing procedure next to the area of interest on the carbon foil.

**Figure 1.**
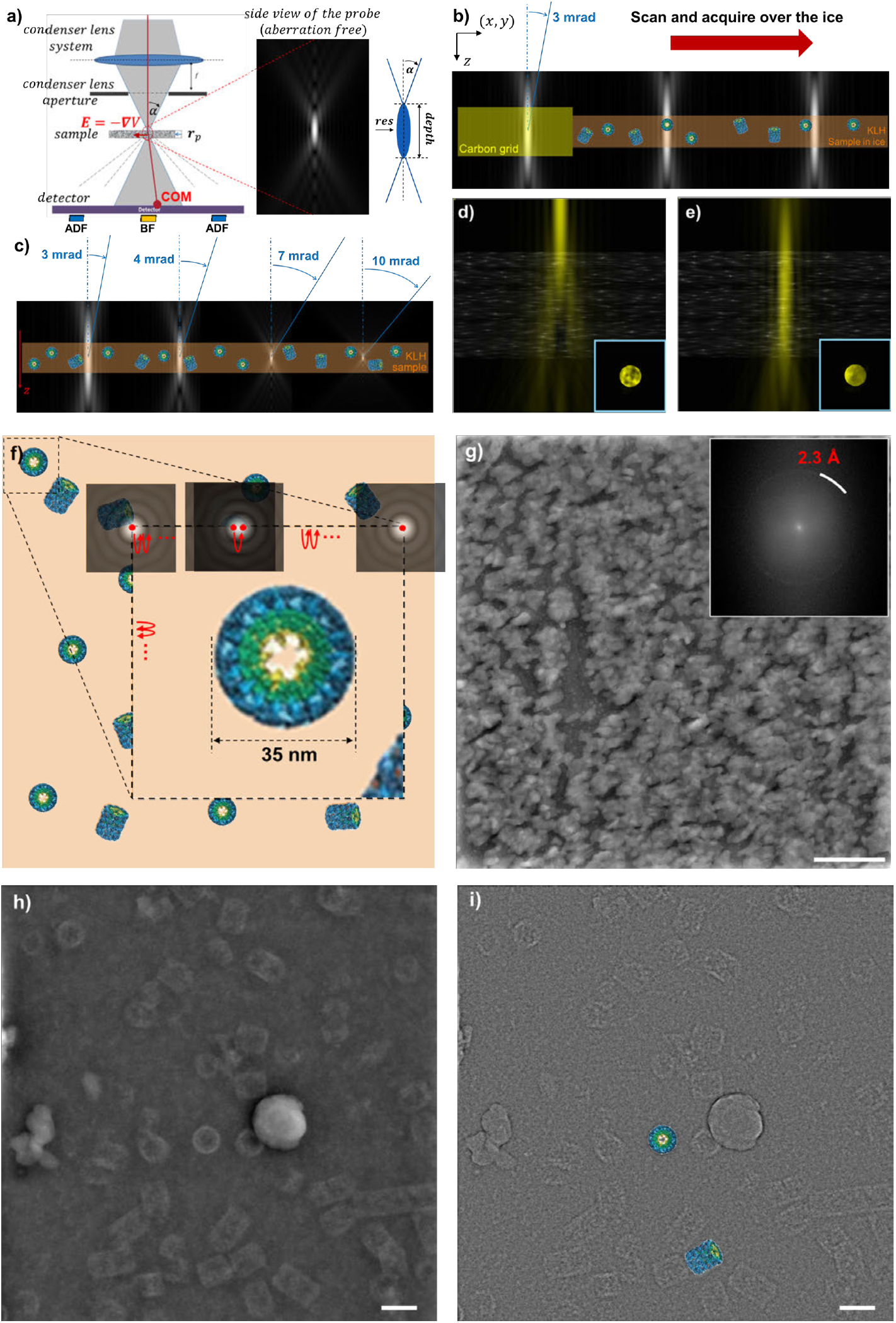
iDPC-STEM imaging setup, acquisition and micrographs. (**a**) Schematic model of the probe formation system, sample and four-quadrant detector. The red line illustrates the average electron path of the beam, the red dot indicates the center of mass (COM) of the intensity at the detector. The center inset (center) corresponds to a longitudinal cross-section through the aberration-free probe. The z-scale along the beam is highly compressed with respect to the xy-plane for visualization (not drawn to scale). On the right, a cartoon model of the probe indicates optical properties: convergence semi-angle (CSA), beam waist width/resolution and length/depth. (**b**) Illustration of the acquisition process: focusing is performed on the carbon foil next to the region of interest (left), before the scanning process starts (center and right). (**c**) Probe shape, i.e., depth of focus, for different CSAs with respect to the sample thickness (not drawn to scale in z direction). (**d**) Focusing using convergence beam electron diffraction (CBED) pattern (inset), out of focus. (**e**) Focusing using CBED pattern (inset), in focus. (**f**) Illustration of the scanning process of typical field of view 512 × 512 nm^2^. The zoomed inset shows step scans using a 0.5 mrad CSA beam as an example. (**g**) Experimental iDPC-STEM micrograph (4096 × 4096) of a gold-on-carbon sample confirming an optical resolution of 2.3 Å (inset top right) acquired at 300 kV, CSA of 4.5 mrad, electron dose of 350 e^-^/Å^2^ and 0.76 Å pixel size, scale bar is 50 nm. (**h**) iDPC-STEM micrograph (4096 × 4096) of vitrified KLH acquired at 300 kV, CSA of 2.0 mrad, with electron dose of 40 e^-^/Å^2^ and 1.5 Å pixel size, scale bar is 50 nm. Note the strong low-frequency contrast of the ice contamination. (**i**) Gaussian high-passed filtered micrograph of (**h**) including atomic models of two different KLH projections drawn to scale.

In order to experimentally verify the optical resolution of the STEM approach, we analyzed a standard 50 nm-thick sample of gold deposited on a carbon film and confirmed the presence of the 2.3 Å gold ring in the power spectrum of the images obtained at 4.5 mrad CSA and electron dose of 350 e^-^/Å^2^ (**Fig. 1g**). The first diffraction ring of the gold lattice is straightforward to detect and the obtained resolution is very close to the theoretical resolution limit of 2.2 Å for the chosen CSA (**Table 1** and **Table S1**). In addition, we note the absence of any typical Thon rings that are common for CTEM imaging (**Fig. S1**). Instead, we observe a four-fold star pattern at the origin of the power spectrum, reflecting the actual CTF of iDPC-STEM. In the ideal case when COM is directly detected, the resulting CTF and power spectrum will be rotationally symmetric and decay towards higher frequencies (**Fig. S2a**). When a four-quadrant detector is used, the resulting two-dimensional CTF has a four-fold pattern in addition to the resolution decay (**Fig. S2b**). The fourfold CTF pattern dominates the very low-resolution frequencies only and has a minor effect on the visual appearance of the image [8, 9] (**Figs. S2c**). The signal further decays towards higher resolutions reaching zero at the theoretical resolution limit of the given CSA (**Fig. S2d and S2e**). Using the described imaging setup, an iDPC-STEM micrograph of ice-embedded keyhole limpet hemocyanin (KLH) was recorded (**Fig. 1h**), which exhibits for instance strong contrast of ice contamination as a prominent low-frequency feature in addition to KLH particles. When the iDPC-STEM micrographs are high-pass filtered, they resemble inverted CTEM images in appearance because of the missing low frequencies (**Fig. 1i**). In conclusion, iDPC-STEM micrographs provide the complete projections of the sample including low and high frequencies [8, 37], which in appearance are comparable to in-focus CTEM images using a phase plate [38].

**Table 1.** Critical STEM imaging parameters for cryo-EM image acquisition at 300 keV electrons (wavelength *λ* = 1.969 pm).

| Convergence semi-angle (CSA) $\alpha$ [mrad] | Experimental STEM resolution based on Au measurement [$\text{\AA}$] | Maximum theoretical STEM resolution $\lambda/(2\alpha)$ [$\text{\AA}$] | Required pixel size for maximum STEM resolution [$\text{\AA}$] | Depth of focus $2\lambda/\alpha^2$ [nm] |
| --- | --- | --- | --- | --- |
| 2.0 | 5.1 | 4.9 | 2.4 | 985 |
| 3.0 | 3.5 | 3.3 | 1.6 | 438 |
| 3.5 | 3.0 | 2.9 | 1.4 | 321 |
| 4.0 | 2.6 | 2.5 | 1.2 | 246 |
| 4.5 | 2.3 | 2.2 | 1.1 | 194 |

In order to quantitatively judge the quality of cryo-iDPC-STEM micrographs, we imaged vitrified TMV as a test specimen. The helical organization of TMV and resulting image repeats are used to assess information transfer of the micrographs. Initial images were acquired with a low CSA of 2.0 mrad at a pixel size of 1.72 Å (**Fig. 2a**). The imaged FOV of 700 × 700 nm is comparable to CTEM cryo-image acquisitions that record a part of a μm carbon hole. The iDPC-STEM micrograph reveals the presence of TMV rods closely packed in rafts close to the edge of the carbon hole. As the contrast results directly from the electrostatic potential, TMV’s protein density is white and not inverted as in CTEM. Closer inspection of the fine structure shows the distances of the 23 Å as well as the 11.5 Å repeats along the helical rod (**Fig. 2b**). Moreover, the Fourier transform shows a continuous information transfer, characteristic TMV layer lines and the absence of any CTF zero crossings (**Fig. 2c**). The noticeable vertical streak along the meridian of the image Fourier transform is a well-known scanning effect in STEM [39]. In images of densely formed TMV rafts using a CSA beam of 4.0 mrad, based on the Fourier transform, we were able to detect high-resolution information up to 1/7.7 and 1/4.6 Å^−1^, which indicated further potential of cryo-iDPC-STEM imaging at high resolution.

**Figure 2.**
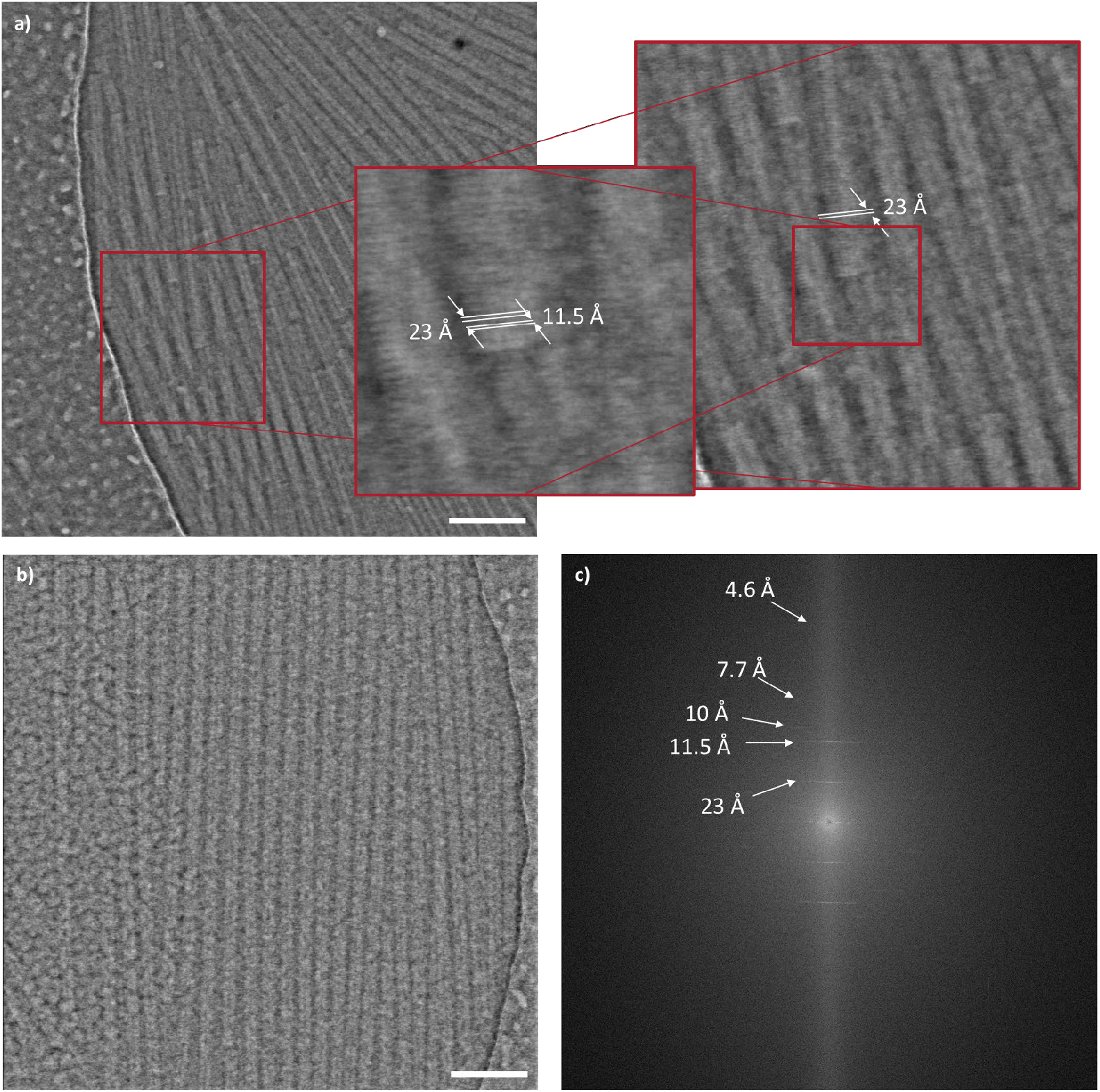
Typical cryo-iDPC-STEM micrographs of tobacco mosaic virus (TMV) acquired at 2.0 and 4.0 mrad convergence semi-angle (CSA). Experimental conditions: voltage 300 kV, pixel size 1.72 Å, image size 4096 × 4096, total electron dose 35 e^-^/Å^2^. (**a**) iDPC-STEM image at 2.0 mrad CSA shows TMV particles in ice with carbon foil on the left, scale bar is 100 nm. Zoomed in regions indicate the helical 11.5 and 23 Å repeats along the rod in real space. (**b**) iDPC-STEM image at 4.0 mrad CSA of a TMV raft, scale bar is 100 nm, and (**c**) the corresponding power spectrum. Layer lines show high-frequency information up to 1/4.6 Å^−1^ (white arrows).

Next, we set out to establish imaging conditions of the TMV sample to obtain high-resolution 3D image reconstructions. As a defocused-based CTEM reference data set, we used a previously determined TMV structure, EMPIAR-10021 [40]. Subsequently, we systematically compared iDPC-STEM data sets taken with beam CSAs of 2.0, 3.0, 3.5, 4.0 and 4.5 mrad respecting critical parameters given in **Table 1**, with the CTEM reference cryo-micrographs. For the detailed comparison of all data sets, we analyzed a typical micrograph including the corresponding Fourier transform (**Fig. 3**). Under all imaging conditions, iDPC-STEM images show strong low-frequency contrast in comparison with the defocused cryo-image. The respective Fourier transforms show the expected 1/23 and 1/11.5 Å^−1^, 1^st^ order and 2^nd^ order layer lines, whose intensities are primarily determined by the number of TMV rods present in the field of view. For more quantitative analyses, we extracted 1300 – 2200 helical segments from several micrographs and computed a power spectrum average, displayed as a 1D helical profile. Comparing the overall slope of the 1D helical profile it is apparent that the profiles derived from the iDPC-STEM images decay faster than from the defocused CTEM micrographs. The relative ratio of the 2^nd^ over 1^st^ order layer line profile peaks increases from 0.2 to 0.5 with CSA beam increase from 2.0 to 4.5 mrad respectively, which indicates improved signal transfer of the higher resolution layer line at higher CSA. The corresponding ratio of the defocused CTEM data set is 0.5 matching the 3.5, 4.0 and 4.5 mrad STEM data sets.

**Figure 3.**
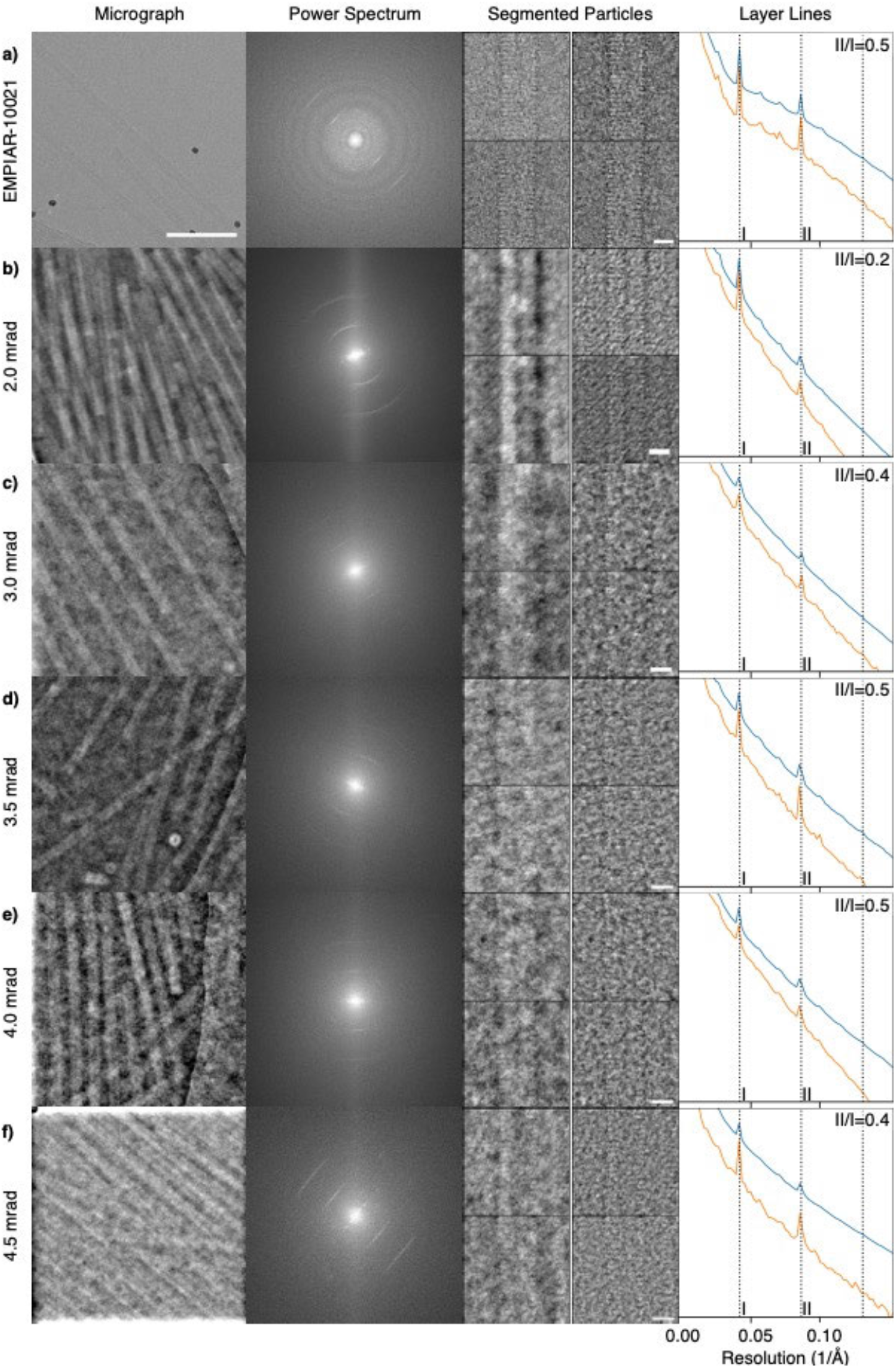
Comparison of TMV cryo-images taken by conventional TEM (CTEM) and iDPC-STEM. Micrograph inset of vitrified TMV (left, scale bar 100 nm), corresponding power spectrum (center left), four in-plane rotated TMV segments (center right: two left column iDPC-STEM segments, two right column high-pass filtered, scale bar 10 nm) and corresponding layer-line profile of added power spectra segments. In the layer line profile, 1^st^ order and 2^nd^ order layer lines give rise to peak I and II at 1/23 and 1/11.5 Å^−1^, respectively. The ratio of II/I is given in the upper right corner. The orange line is averaged from the corresponding data sets of around 2000 segments and the blue line is obtained from averaged segments of a single micrograph. (**a**) CTEM (defocused-based) cryo-images as well as (**b**) 2.0 mrad, (**c**) 3.0 mrad, (**d**) 3.5 mrad, (**e**) 4.0 mrad and (**f**) 4.5 mrad beam CSA iDPC-STEM cryo-images.

Finally, using the individual CTEM and iDPC-STEM micrographs, we processed approximately the same number of helical segments from the different data sets using the typical single-particle helical reconstruction work-flow of RELION [41] (**Table 2**). It should be noted, however, that for iDPC-STEM in the absence of defocusing, we neither used CTF determination nor any CTF correction options. Subsequently, we computed the cryo-EM 3D TMV structures using the 2.0, 3.0, 3.5, 4.0 and 4.5 mrad beam-CSA data sets and determined the TMV structures at 6.3, 4.3, 3.9, 3.5 and 3.7 Å resolution, respectively, based on the 0.143 Fourier shell correlation (FSC) cutoff criterion [42, 43] (**Fig. S3a**). The trend of near-atomic resolution values of high CSA acquisitions at 3.5, 4.0 and 4.5 mrad could also be independently confirmed by a mask-less FDR-FSC determination approach [44] at 3.4, 3.2 and 3.3 Å, respectively (**Fig. S3b**). The reprocessed CTEM data set went to slightly poorer resolution of 3.7 and 3.5 Å for the 0.143 and the FDR-FSC cutoffs, respectively. Local resolution assessment corroborates the better map quality of the 4.0 mrad CSA iDPC-STEM structure in comparison with the CTEM structure (**Fig. S3c**).

**Table 2.** Summary of 3D reconstruction results of reference CTEM and described cryo iDPC-STEM data sets.

|  | CTEM/EMPIAR-10021 | iDPC-STEM 2.0 mrad | iDPC-STEM 3.0 mrad | iDPC-STEM 3.5 mrad | iDPC-STEM 4.0 mrad | iDPC-STEM 4.5 mrad |
| --- | --- | --- | --- | --- | --- | --- |
| Pixel size [$\text{\AA}$] | 1.062 | 1.705 | 1.23 | 0.98 | 0.98 | 0.75 |
| Number of segments | 2068 | 2073 | 1392 | 1330 | 2224 | 1678 |
| Number of asymmetric units | 53,768 | 87,066 | 41,760 | 31,920 | 75,616 | 57,052 |
| B-Factor [$\text{\AA}^2$] | 126 | 492 | 155 | 105 | 132 | 126 |
| Maximum resolution target (Table 1) [$\text{\AA}$] | N/A | 5.1 | 3.5 | 3.0 | 2.6 | 2.3 |
| Resolution expected range (Table S1) [$\text{\AA}$] | N/A | [7.4 - 5.6] | [5.0 - 3.8] | [4.3 - 3.3] | [3.8 - 2.9] | [3.3 - 2.5] |
| Resolution FSC (0.143<sup>[42]</sup>/ FDR<sup>[44]</sup>) [$\text{\AA}$] | 3.8 / 3.6 | 6.3 / 5.9 | 4.3 / 4.1 | 3.9 / 3.4 | 3.5 / 3.2 | 3.7 / 3.3 |

To validate the numerical resolution assessment, we inspected the CTEM and iDPC-STEM cryo-EM density maps in more detail (**Fig. 4a**). In the presence of a previously refined atomic model (PDB-ID 4UDV) [40] we find that secondary structures are well discernible for the 2.0 mrad map at 6.3 Å resolution (**Fig. 4b**). The density of the α-helical pitch can be well recognized for the 3.0 mrad map at 4.3 Å resolution (**Fig. 4c**). In addition, the maps also show clear density of the RNA and for ribose moieties that are tightly packed between the subunits. Qualitatively, the molecular features that are discernible in 3.5, 4.0 and 4.5 mrad maps at near-atomic resolution constitute larger side aromatic and positively charged chains such as F35, R41, W52, F62, F87, R113 and R122 in addition to a well-defined polypeptide backbone (**Fig. 4d-f**). In analogy to the CTEM map that received the same total electron dose of 35 e^-^/Å^2^, density for negatively charged side chains D77, E106, D115, D116 and E131 is largely absent. The displayed cryo-EM densities were sharpened based on the Guinier plot of the 3D reconstruction. The corresponding B-factors were determined at 492, 155, 105, 132 and 126 Å^2^ for the 2.0, 3.0, 3.5, 4.0 and 4.5 mrad maps, respectively. The smaller B-factors correlate with the generally improved resolution and image quality for the 3.5 – 4.5 mrad data sets. Together, the cryo-EM density analysis of vitrified TMV confirms that in-focus iDPC-STEM images can be reliably used to resolve the detailed protein structures of ice-embedded biological macromolecules to near-atomic resolution.

**Figure 4.**
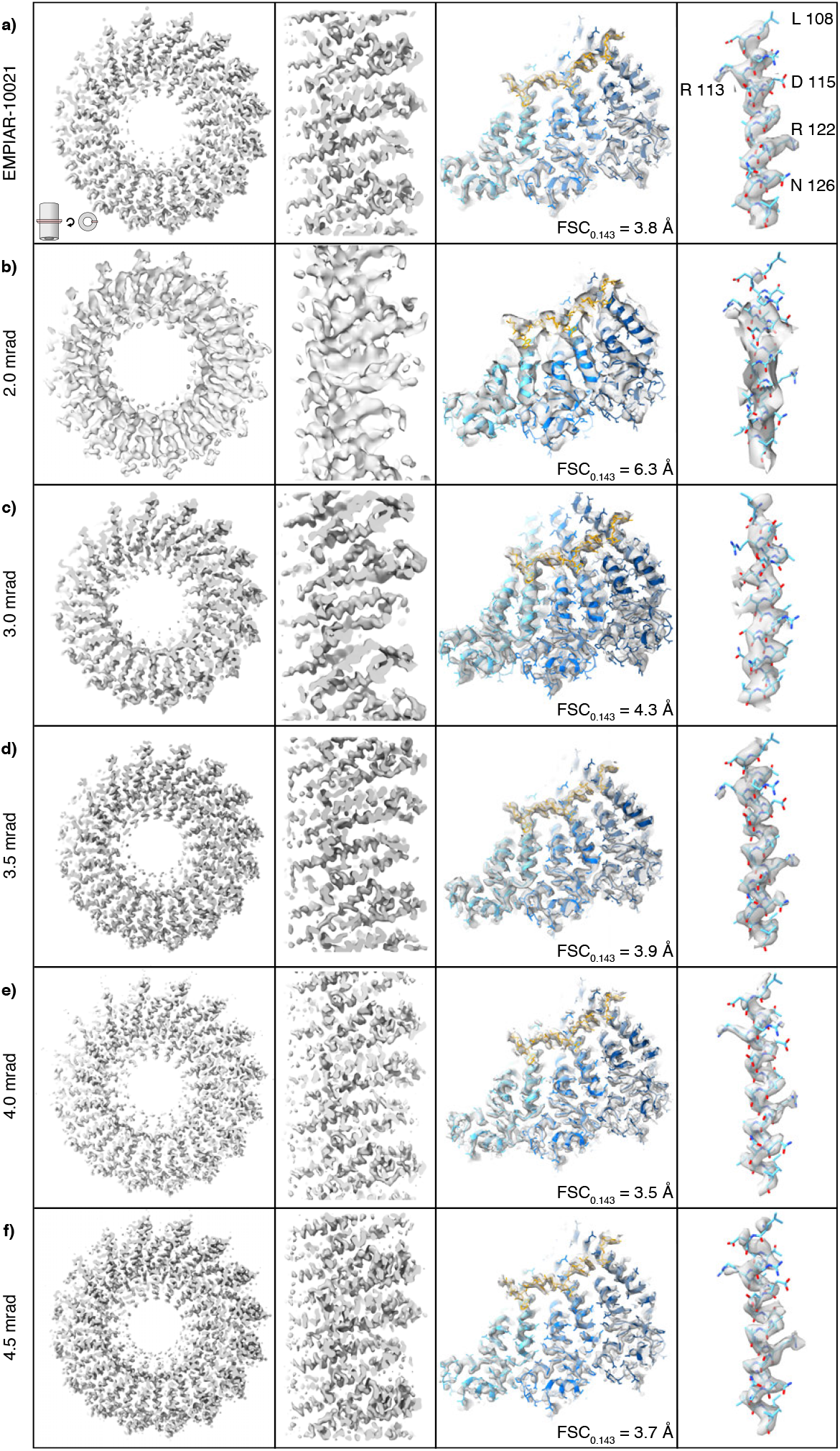
Comparison of cryo-EM density maps reconstructed from conventional TEM (CTEM) EMPIAR-10021/PDB ID 6SAG and iDPC-STEM images using different convergence semi-angle (CSA) beams. Top slab view of TMV map (left), enlarged side slab view of multiple TMV subunit densities (center left), three-subunit top view (center right) and detailed side-chain density of α-helix (108-130). Cryo-EM densities from (**a**) CTEM, (**b**) 2.0 mrad, (**c**) 3.0 mrad, (**d**) 3.5 mrad, (**e**) 4.0 mrad and (**f**) 4.5 mrad beam CSA iDPC-STEM images. The EMPIAR-10021 CTEM map has a resolution of 3.8 Å and the 2.0 / 3.0 mrad iDPC-STEM maps at 6.5 / 4.3 Å resolution, respectively. The 3.5 mrad, 4.0 mrad and 4.5 mrad CSA cryo-iDPC-STEM maps share the same molecular features with a CTEM 3.8 Å near-atomic resolution map. The 4.0 mrad CSA iDPC-STEM data set gives the best resolution of 3.5 Å of compared data sets. All near-atomic resolution maps lack density for negatively charged residues (e.g. D115).

## Discussion

Using iDPC-STEM, we imaged plunge-frozen TMV as a biological test specimen, assessed the image quality and analyzed the resulting 3D cryo-EM density maps. We demonstrated that iDPC-STEM produces high-contrast micrographs displaying the regular helical image features of TMV. When we subjected multiple micrographs to the single-particle helical reconstruction workflow, we determined several near-atomic resolution structures using different iDPC-STEM imaging conditions down to 3.5 Å. By using the appropriate combination of CSA beam and associated imaging parameters we demonstrated the capability of obtaining resolution within the expected theoretical limits (**Table 2**). These data establish that iDPC-STEM imaging of cryo-vitrified biological samples generates micrographs of sufficiently high quality suitable for near-atomic resolution single-particle cryo-EM structure determination.

One of the apparent features of the recorded iDPC-STEM cryo-micrographs is that they show continuously transmitted contrast over the complete frequency band. This favorable property is specific to the iDPC-STEM approach. For instance, annular bright field (ABF)-STEM suffers from CTF shortcomings similar to CTEM. Due to the reciprocity theorem ABF-STEM also requires defocus to generate contrast and to enhance the low-frequency transfer [45]. In addition, only part of the scattered electrons is collected by the detector and used to form an image. For ADF-STEM, imaging can be performed in focus resulting in an overall positive CTF. However, the total number of electrons collected in the dark field is several orders of magnitude smaller than in the bright field part, making ADF detection very dose-inefficient [14]. Moreover, for ADF-STEM the imaged object corresponds to the square of the electrostatic potential [8, 45], causing the light elements to disappear when imaged next to heavier ones. Based on these considerations, standard STEM techniques do not have sufficiently good properties to be exploited for low-dose imaging of radiation-sensitive materials. Conversely, iDPC-STEM uses all the available electrons, has a favorable CTF devoid of any contrast reversal and CTF zero-crossings. The imaged object is directly proportional to the electrostatic potential field of the sample [8, 45]. When imaged gold particles deposited on a carbon film using iDPC-STEM with a 4.5 mrad CSA beam, we resolved the details to 2.3 Å resolution, confirmed by the first gold-lattice diffraction ring in the power spectrum of the image (**Fig. 1**). Due to the detector architecture of four quadrants, we also observed a four-fold shaped 2D CTF. This CTF pattern can be compensated by a 2D CTF correction [9] as the theory is well understood [7, 8]. However, we did not find this necessary in our analysis, as helical segments with random orientations within the ice layer plane are averaged. The power spectra of the recorded iDPC-STEM cryo-micrographs did not show any Thon rings that are common for defocused-based CTEM imaging. The resulting high-resolution signal transfer in iDPC-STEM is, therefore, very well suited for near-atomic cryo-EM structure determination.

The performance of iDPC-STEM depends on experimental parameters that must be controlled during imaging. The beam-CSA critically determines the maximum possible resolution of the STEM image (**Table 1**). This resolution is exactly related to the beam width and the beam depth of focus, also referred to as depth resolution or beam-waist length, according to theoretical considerations [45, 46, 47]. When increasing the CSA, higher resolution is obtained provided the probe is aberration free while the beam depth of focus becomes smaller. Consequently, when sample thickness exceeds the decreasing beam-waist length, the images represent optical depth sections rather than projections [45, 48]. We estimate that the sample regions imaged here have thicknesses of 18 to 54 nm corresponding to a diameter of a single TMV rod and three TMV rods, respectively. When typical ice thicknesses of 10 - 60 nm, common for plunge-frozen specimens, are considered, only the beam-CSAs larger than 9 mrad at 300 kV will not be sufficient for obtaining complete projections of the sample. With the maximum beam-CSA of 4.5 mrad used in this study resulting in a depth of focus of 194 nm, complete projections are produced in all cases. For higher CSAs, e.g. larger than 7.0 mrad, in addition to decreasing the depth of focus, the spherical aberration of the probe deteriorates the image due to the introduction of CTF zero crossings [47]. This property justifies the usage of C_s_-correctors when higher spatial resolutions are targeted. The discussed beam relationship of resolution vs. depth of focus also opens up principal possibilities for high-resolution imaging of thicker cryo-frozen samples using optical sectioning STEM [37, 49].

The estimated resolutions of the determined iDPC cryo-EM structures confirm the basic optical considerations that increasing CSA leads to higher spatial resolution. In order to evaluate additional limiting parameters of the imaging setup, we simulated iDPC-STEM images of a hemoglobin molecule embedded in vitreous ice. First, upon increasing the CSA beam, we observed a decrease in the signal-to-noise (SNR) of the particle (**Fig. S4a**). When the same number of electrons is used, smaller CSA beams will produce a higher SNRs due to the reduced solid-angle volume of the electron-wave behind the sample. Therefore, for initial micrograph assessment, smaller beam-CSAs are recommended for boosting SNR. The apparent SNR can also be matched by increasing the electron dose at the expense of radiation damage effects. Second, at high CSAs and large pixel sizes, aliasing may obstruct the image. To illustrate this effect in simulated images, we employed a large CSA beam of 10 mrad at a pixel size larger than the resolution cutoff imposed by the CSA. Under these conditions, the discernible particle contrast weakened because higher resolution information is folded back to low frequencies (**Fig. S4b**). After we applied a pixel size smaller than the CSA cutoff, the effect was not present. This form of aliasing is particularly relevant in the case of biological cryo-samples where the surrounding amorphous ice provides additional high-frequency signal. To avoid aliasing, we routinely adjusted the scan interval, i.e., pixel size, such that the frequency cutoff given by beam-CSA is smaller than the expected Nyquist frequency (**Table 1**). For example, the highest resolution iDPC-STEM TMV map at 3.5 Å resolution was achieved with the 4.0 mrad CSA beam and a pixel size of 0.98 Å using an electron dose of 35 e^-^/Å^2^.

The re-processing of a comparable TMV CTEM data set (EMPIAR-10021) recorded on a Falcon II DED camera [40] of approximately 50,000 asymmetric units and a deposited electron dose of 35 e^-^/Å^2^ resulted in a slightly poorer resolution map of 3.7 Å. The 4.0 mrad iDPC-STEM map showed the expected cryo-EM density features at the given resolution (**Fig. 4**). The map sharpening B-factors determined by Guinier analysis of both the CTEM and 4.0 mrad iDPC-STEM maps are at around 130 Å^2^. For poorer resolutions of TMV data sets taken on film, higher B-factors of 240 and 280 Å^2^ were reported [29, 52] whereas for TMV data sets taken on the 3^rd^ generation of Falcon III DED and K2 cameras, lower B-factors of 100 and 40 Å^2^ have been determined [33, 53] (**Table 3**). Therefore, it is remarkable that iDPC-STEM, in combination with a four-quadrant detector without any motion correction, shows a very similar performance and a B-factor as in CTEM imaging with a 2^nd^ generation DED camera including motion correction. Although the map features indicate that the final iDPC-STEM map exhibits very similar radiation damage effects on the negatively charged side chains as other CTEM maps [40, 50, 51], the iDPC-STEM spot-scanning approach appears not to suffer critically from beam-induced movement that used to be one of the critical resolution-deteriorating issues before the introduction of movie mode in DED cameras [54].

**Table 3.**
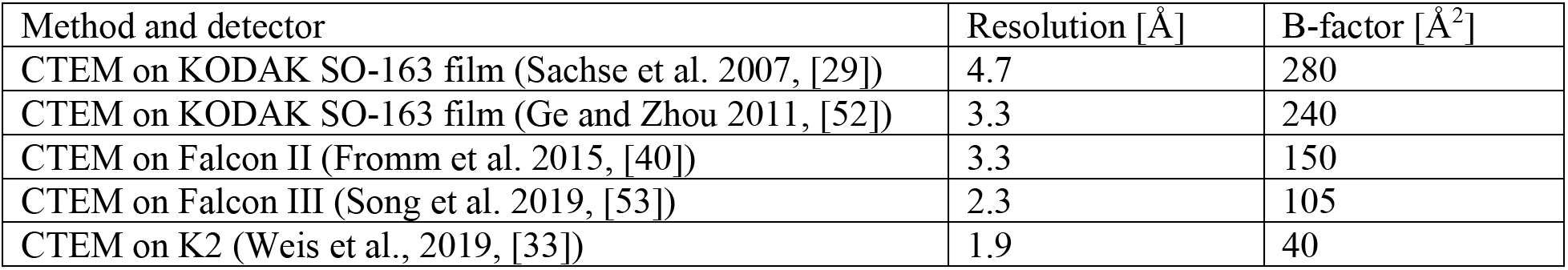
Reported TMV structures based on different CTEM data sets with reported resolution and B-factors.

| Method and detector | Resolution [$\text{\AA}$] | B-factor [$\text{\AA}^2$] |
| --- | --- | --- |
| CTEM on KODAK SO-163 film (Sachse et al. 2007, [29]) | 4.7 | 280 |
| CTEM on KODAK SO-163 film (Ge and Zhou 2011, [52]) | 3.3 | 240 |
| CTEM on Falcon II (Fromm et al. 2015, [40]) | 3.3 | 150 |
| CTEM on Falcon III (Song et al. 2019, [53]) | 2.3 | 105 |
| CTEM on K2 (Weis et al., 2019, [33]) | 1.9 | 40 |

Working out the appropriate iDPC-STEM parameters for imaging of vitrified specimens was straightforward as TMV required relatively few micrographs for quantitative image analysis and 3D reconstruction. The scanning procedures used for the micrograph acquisition took approximately 20 – 80 seconds for the 5.0 mrad and the 2.0 mrad CSA beams, respectively **(Table 4)**. This way, approximately 60 micrographs were collected in each CSA session in the absence of any automation. Collecting large data sets up to 300 micrographs per hour common for typical CTEM single-particle acquisitions would not be possible in this manner [55]. Yet, automation of the STEM acquisition procedures will be straightforward to employ in the future. Detectors used for iDPC-STEM allow scan speeds of two orders of magnitude faster than the one used in this work, which will ultimately reduce acquisition times to below one second. This way, when automation is further implemented, it will become possible to image equally large data sets of single-particle specimens in similar time frames as in CTEM. Additionally, recording individual frames in movie mode will reduce the effects of beam-induced motion and further improve the image quality of cryo-iDPC-STEM. Once iDPC-STEM imaging can be applied to tilt series of thicker biological specimens, cryo-electron tomographic reconstructions will benefit from the complete contrast transfer in STEM images devoid of any oscillating CTF while still preserving high-resolution information. Moreover, alternative STEM acquisition schemes and micrograph reconstruction approaches should also prove beneficial for imaging biological samples [26]. The results presented here, using iDPC-STEM to image vitrified TMV specimens, reveal that STEM approaches should be further explored for improved high-resolution cryo-EM structure determination.

**Table 4.** Scanning times with a total electron dose of 35 e^-^/Å^2^ using 4 pA beam current.

|  |  |  |  |  |  |
| --- | --- | --- | --- | --- | --- |
| CSA [mrad] | 2.0 | 3.0 | 3.5 | 4.0 | 4.5 |
| Pixel size [$\text{\AA}$] | 1.70 | 1.23 | 0.98 | 0.98 | 0.75 |
| Dwell time [$\mu\text{s}$] | 4.05 | 2.12 | 1.35 | 1.35 | 0.79 |
| Frame time for 4096 x 4096 image [s] | 67.9 | 35.6 | 22.6 | 22.6 | 13.2 |

## Materials and Methods

### Specimen preparation

Quantifoil grids were rendered hydrophilic using a glow discharge device (GloQube Plus, Quorum) for 45 seconds in air before specimen application to the grid. An aliquot of 3.5 μl stock solution of tobacco mosaic virus (TMV) at a concentration of approximately 90 mg/ml or 2.5 μMol was applied onto a 200 mesh Quantifoil grid with regular R2/2 hole carbon support. The excess of the applied droplet was blotted using a Vitrobot Mark IV (Thermo Fisher Scientific MSD). Due to the high concentration, optimal results for obtaining thin ice across the 2 μm holes were achieved using a high blot force of +10 and a duration for blotting of 10 seconds, prior to plunge freezing. Grids were prepared at 4 °C and 100% humidity. To maximize occupancy of holes with rafts of TMV rods, application of TMV at high concentrations was critical. After flash freezing, grids were stored in grid boxes in liquid nitrogen for subsequent mounting into autogrids of the Krios G4 Cryo-Autoloader. For the keyhole limpet hemocyanin (KLH) samples, cryo-grids were prepared as described above. For KLH, however, a 10 mg/ml or 2.5 μMol solution was plunge frozen using a blot force of −10 and 6 seconds blotting time.

### Scanning transmission electron microscopy (STEM)

STEM imaging was conducted using a Thermo Fisher Scientific Titan Krios G4, operated at 300 kV. The column was equipped with a standard high-brightness field emission gun (X-FEG), three-condenser lens system, C-TWIN objective lens with wide-gap pole piece (11 mm and C_s_ = 2.7 mm), Panther segmented STEM detector, a Ceta camera and Falcon 4 direct electron detection (DED) camera. A combination of different C2 apertures (20 μm and 50 μm) and C2/C3 lens current ratios were used to create different CSA of the beam. Alignment for STEM was done by aligning beam shift, beam tilt pivot points and beam tilt in STEM mode for the different convergence angles. For accurate determination of the COM, a de-scan alignment has additionally been performed. The CSA of the applied beams were measured with high precision using the Au cross-grating and Ceta camera by first recording the radii of the gold diffraction rings for calibration and subsequently measuring the radius of the bright-field (BF) disc. The beam current was determined using the fluorescent (flu-) screen, calibrated by a Faraday-cup measurement. The total electron dose delivered per unit square area is given by

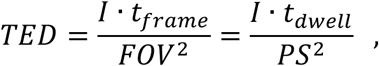

where *I* is the beam current (number of electrons per unit time) while *t_frame_* is total scanning time needed to scan the whole field of view *FOV* frame area. Further, *t_dwell_* is the time that the beam spends per pixel, and *PS* is the pixel size. Note that *t_frame_* = *n* · *t_dwell_* where, *n* = *N*^2^ number of pixels of e.g. *NxN* size image, while *FOV* = *N* · *PS*. Therefore, the number of pixels cancels and yields the right-hand side of the equation. The beam current was held at fixed 4 pA while the dwell time was used as a parameter to fine-tune the final total dose of 35e^-^/Å^2^. Low magnification atlas screening, in searching for suitable ice thickness area, was performed in TEM mode with the Falcon 4 DED and the MAPS software. Intermediate magnification ADF-STEM images were collected at 5000 times magnification to get a STEM overview image and assign it to the TEM atlas. Using the MAPS software, the different magnifications and modes were correlated and aligned. A low magnification atlas (e.g. a tiled set of images representing most of the grid area), served as a navigation map for finding grid squares that have thin ice and sufficiently dense rafts of TMV specimen. Smaller 4 × 4 or 5 × 5 tiled atlas maps were created within the grid squares to navigate the stage to the holes containing TMV. Data was acquired and stored at each tile position across the holes using Velox software.

### iDPC-STEM imaging and acquisition

For iDPC-STEM acquisition, the four-quadrant configuration of the Panther STEM detection system (Thermo Fisher Scientific) was used. For every convergence angle, the camera length was chosen such that bright-field disk of the beam at the detector covers detector area with a four times larger radius than the radius of the central hole. For each CSA beam ≥ 4.5mrad, gold rings were used in the power spectrum of STEM (iDPC, **Fig. 1c** and **Fig. S2c,d** and ADF, **Fig. S1**) to confirm the resolution by imaging a standard cross grating gold-on-carbon sample. As mentioned above, the total applied electron dose 35 e^-^/Å^2^ was used according to **Table 4**. Therefore, the typical acquisition time for a 4096 × 4096 sized micrograph was between 13.2 seconds (smallest, 0.75Å pixel size) and 67.9 seconds (largest, 1.7Å pixel size). Approximately 10 s rest-time was applied to the stage between acquisitions. Focusing has been performed at the carbon grid next to the area of interest (hole with ice and particles) by judging the flatness of the CBED pattern (Ronchigram). Typically, 50 to 70 micrographs have been acquired per CSA session. The micrographs with the highest resolving layer lines (approx. one third) were selected and used for helix segmentation and 3D reconstruction. For smooth data recording, the procedure was followed in MAPS software (ThermoFisher Scientific) for optimal, accurate and safe stage navigation. The overview maps served to navigate and collect high-resolution iDPC-STEM images preventing multiple exposures.

### Image processing and 3D single-particle helical reconstruction

The raw iDPC-STEM micrographs were Gaussian high-pass filtered with a FWHM of 251 Å. Helical coordinates were interactively picked using EMAN2 [56]. The in-plane rotated segments were used to calculate the average power spectra. The power spectra sums were collapsed in the direction orthogonal to the helical axis into 1D spectra as described [57]. Collapsed power spectra were used to calibrate the pixel size by matching the visible layer lines with the expected layer lines of 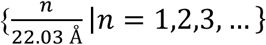. In the case of the 2.0 mrad CSA data set, a varying pixel size was detected and therefore, two out of four micrographs were rescaled by max. 1.8 % to the smallest determined pixel size. Rescaling was not necessary for any other CSA data set. Further image processing was performed using Relion 3.1 [58]. Depending on the data set, one or two rounds of 2D classifications with 10 classes were performed. For the following steps, only particles from classes showing high-resolution details of TMV were included. Due to the absence of defocusing in iDPC-STEM and overall positive CTF [8, 9], neither CTF determination nor any CTF correction option was used. The EMPIAR-10021 CTEM data set was re-processed according to the standard single-particle helical reconstruction workflow. To ensure comparability with the iDPC-STEM data sets, motion correction was performed limiting the included frames to a total dose of 35 e^-^/Å^2^ with the exposure/dose weighting option turned off. To generate a reference for the 3D refinement, 3D classification with one class and a featureless cylinder as a reference was performed. After refinement, a mask was created from one of the half maps, including the central 30 volume % in z-direction. Subsequently, post-processing and local resolution estimation were performed. The half maps were used for FSC calculation including mask deconvolution [43] and resolution taken at the 0.143 threshold [42]. To validate the estimated resolutions with another criterion, the mask-free FDR-FSC approach [44] was applied to the central 30 volume % in z-direction. Cryo-EM density analysis was performed using available TMV coordinates PDB-ID 4UDV [40] and 6SAG [33]. The atomic models were rigid-body fitted in the map, displayed and Figures prepared using UCSF Chimera, UCSF ChimeraX and Coot [59, 60, 61].

### iDPC-STEM imaging simulations

All STEM image simulations (applied in **Fig. 1d** and **e**, and in **Fig. S4**) were produced using the multi-slice method [47], extended to support the iDPC-STEM as explained in the following work on several applications [7, 8, 37, 45]. The parameters for simulations were chosen to accommodate conditions given in **Fig. S4**. Noise was added at the quadrant detection level based on electron dose used in the experiments and varied to illustrate the effect on the image.

## Acknowledgments

The authors would like to thank the following people for valuable support, remarks and discussions during conducting the experiments, performing 3D reconstructions and/or preparing this manuscript: Thomas Hoffmann, Emrah Yücelen, Oliver F. Raschdorf, Sorin Lazar, Yuchen Deng, Roland Jonkers, Bas Groen, Arno Meingast, Bert de Vries and Stefano Vespucci. The authors gratefully acknowledge the computing time granted through JARA on the supercomputer JURECA at Forschungszentrum Jülich [62]. K. M.-C. and M.-L. L. acknowledge funding from the Helmholtz society under contracts VH-NG-1317 (moreSTEM) and ZT-I-0025 (Ptychography 4.0).

## Availability of data

All data needed to evaluate the conclusions in the paper are presented in the paper and/or the Supplementary Materials. The EMDB accession numbers for the reconstructed TMV cryo-EM maps taken at different CSAs are EMD-xxx (CSA:2.0), EMD-xxx (CSA: 3.0), EMD-xxx (CSA: 3.5), EMD-xxx (CSA: 4.0) and EMD-xxx (CSA: 4.5).

## Author contributions

I.L. and C.S. designed the research. F.d.H. prepared TMV cryo-specimen. M.W. optimized the STEM column alignment, adjusted imaging conditions for low dose imaging and designed the acquisition workflow. M.W. and I.L. operated electron microscope in iDPC-STEM mode under these conditions. Initial data evaluation was performed by I.L., R.E. and E.V.P. Supporting simulations were performed by E.G.T.B. Image processing and single-particle helical reconstruction was done by M.B., M.L.L., K.M.K. and C.S. The manuscript was written by I.L., M.L.L. and C.S. with input from all the authors.

## Supplementary Material

### Supplementary Tables

**Table S1.** Extended imaging parameters at 300kV (wavelength *λ* =1.969 pm and condenser system spherical aberration *C_S_* = 2.7 mm). Related to Table 1.

| Convergence semi-angle (CSA) $\alpha$ [mrad] | Experimental STEM resolution based on Au measurement [$\text{\AA}$] | Maximal STEM resolution $\lambda/(2\alpha)$ [$\text{\AA}$] | Required pixel size at maximum STEM resolution [$\text{\AA}$] | Depth of focus $2\lambda/\alpha^2$ [nm] |
| --- | --- | --- | --- | --- |
| 2.0 | 5.6 | 4.9 | 2.4 | 985 |
| 3.0 | 3.8 | 3.3 | 1.6 | 438 |
| 3.5 | 3.3 | 2.9 | 1.4 | 321 |
| 4.0 | 2.9 | 2.5 | 1.2 | 246 |
| 4.5 | 2.5 | 2.2 | 1.1 | 194 |
| 5.0 | 2.3 | 2.0 | 1.0 | 158 |
| 6.0 | 1.8 | 1.6 | 0.8 | 109 |
| 7.0 | 1.6 | 1.4 | 0.7 | 80 |
| Values above are valid for spherical aberration non-corrected probes up to $C_S = 2.7$ mm | | | | |
| Critical maximal convergence semi-angle after which probe deteriorates rapidly in all three directions is given with $\alpha_{max} = 1.34 \lambda / (C_S \lambda^3)^{1/4}$, according to theoretical considerations ([46], Sec. 3.5.1) | | | | |
| $\alpha_{max} = 7.0$ mrad | | | | |
| Values below are well-defined and valid for aberration corrected probe |  |  |  |  |
| 8 | 1.4 | 1.2 | 0.6 | 62 |
| 9 | 1.3 | 1.1 | 0.5 | 49 |
| 10 | 1.2 | 1.0 | 0.5 | 39 |
| 20 | 0.6 | 0.5 | 0.2 | 10 |
| 30 | 0.4 | 0.3 | 0.1 | 4 |

### Supplementary Figures

**Figure S1.**
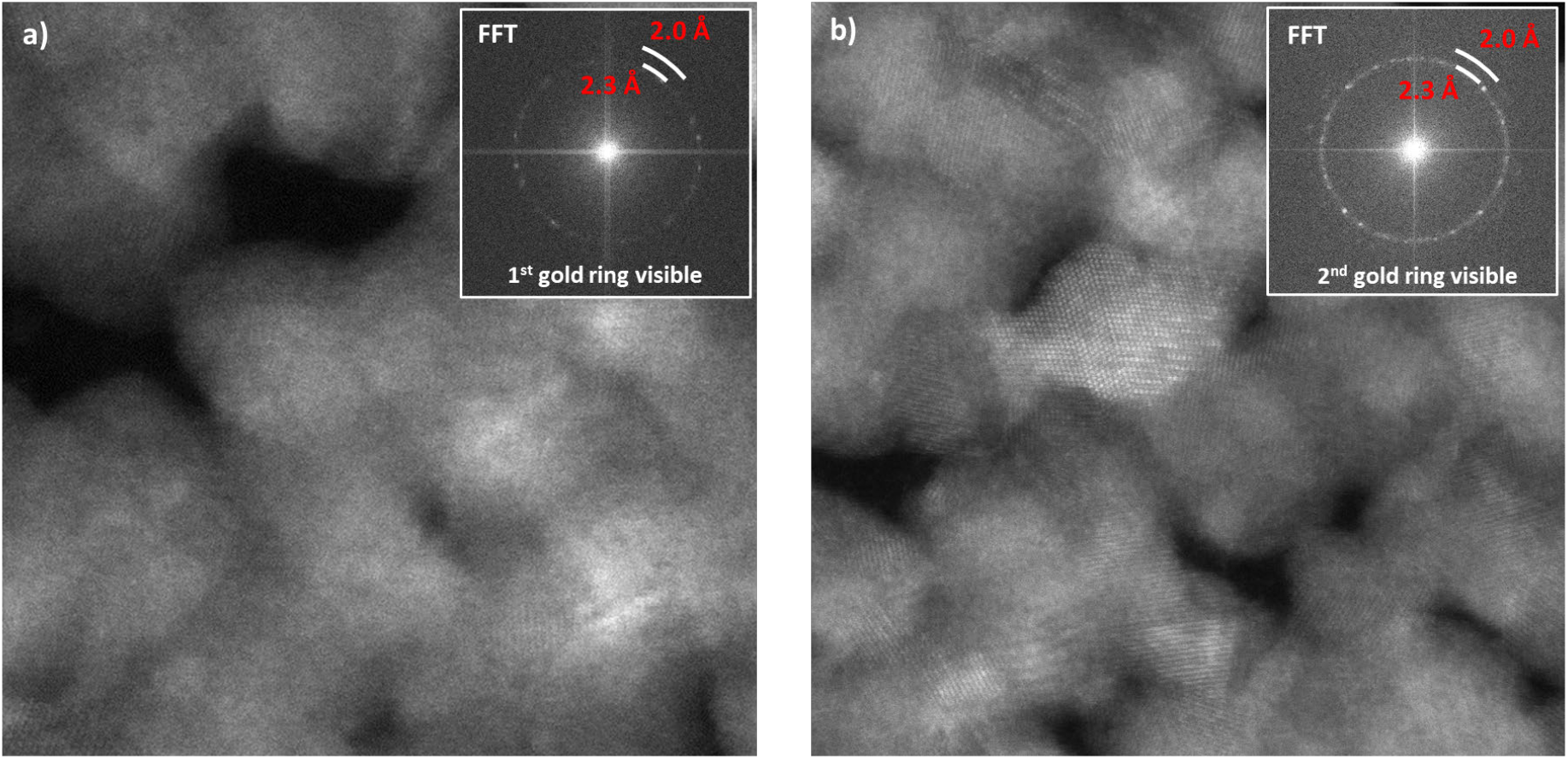
Standard experimental resolution test on gold grating using ADF-STEM. (**a**) ADF-STEM image acquired with 5.0 mrad CSA beam. Power spectrum of the image shows the first gold ring corresponding to a resolution of 2.3 Å. The second gold ring is not visible at a 5.0 mrad CSA beam due to the CTF. (**b**) The same as (**a**) taken with CSA of 6.0 mrad, showing the presence of a second gold ring at a resolution of 2.0 Å. High spatial STEM resolution is confirmed on standardized gold-on-carbon sample for high-dose imaging conditions (>10^4^ e^-^/Å^2^).

**Figure S2.**
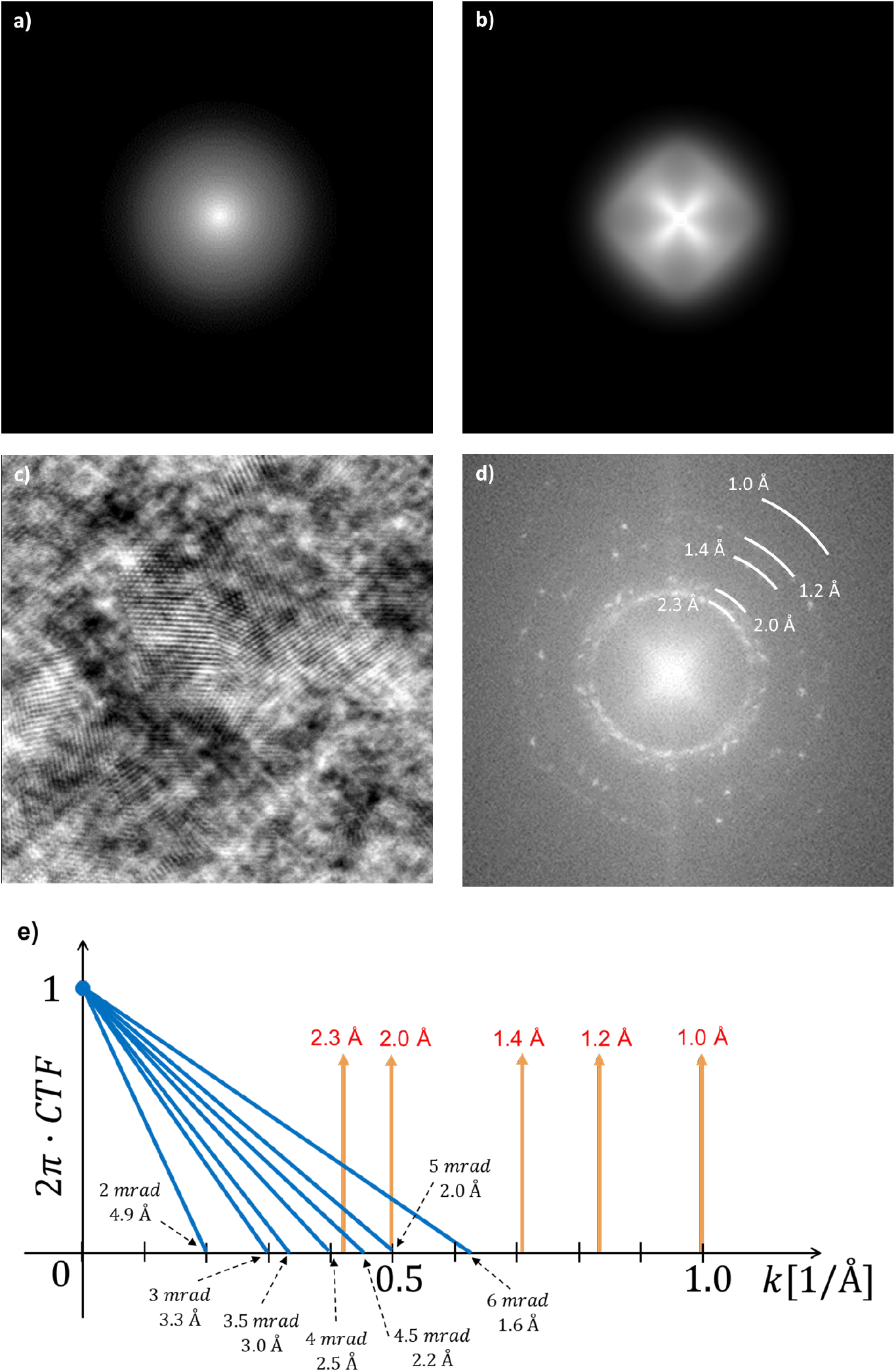
Theoretical 2D CTF’s of ideal integrated center of mass (iCOM)-STEM vs. iDPC-STEM using four-quadrant detector. (**a**) Theoretical 2D rotationally symmetric CTF of iDPC-STEM based on an ideal COM detector (iCOM-STEM). (**b**) Theoretical 2D CTF of the iDPC-STEM using a four-quadrant detector, reflecting the fourfold symmetry of the detector [7, 8]. (**c**) Example of iDPC-STEM image of gold on amorphous carbon sample using a four-quadrant detector at 300 kV (CSA: 20 mrad, C_s_-corrected, dose: 10^4^ e^-^/Å^2^). (**d**) Power spectrum of iDPC-STEM with characteristic gold rings of corresponding gold planes (distances indicated). Note, the four-fold cross reflection of the CTF in the center and the absence of Thon rings. (**e**) Simplified azimuthally averaged CTF profiles (illustrated as straight lines, for the exact shape see [7, 8]) of all CSA beams used in this work with respect to gold ring positions in reciprocal space (yellow arrows).

**Figure S3.**
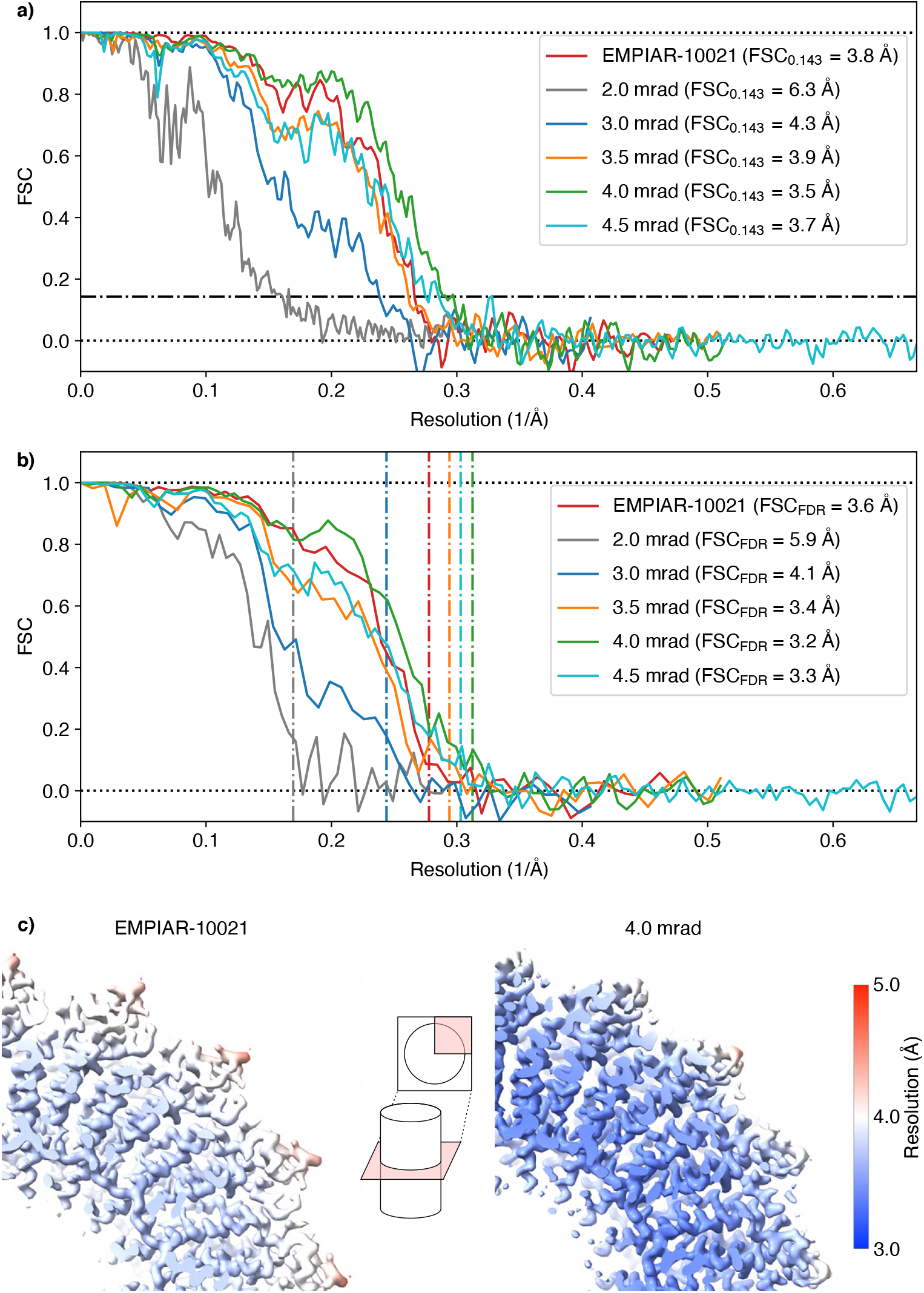
Fourier shell correlations (FSC) used for resolution estimation of CTEM and iDPC-STEM data sets of different CSA beams including local resolution assessment. (**a**) Resolution estimation by FSC including mask deconvolution using the 0.143 criterion [43]. (**b**) Resolution estimation by the mask-free FDR-FSC criterion [44]. (**c**) Color-coded local resolution estimates based on FDR-FSC [44] superimposed on molecular density of CTEM EMPIAR-10021 structure (left) and iDPC-STEM structure (right) acquired using CSA of 4.0 mrad. The best resolution of the compared cryo-EM maps is obtained for the iDPC-STEM structure acquired at 4.0 mrad CSA beam.

**Figure S4.**
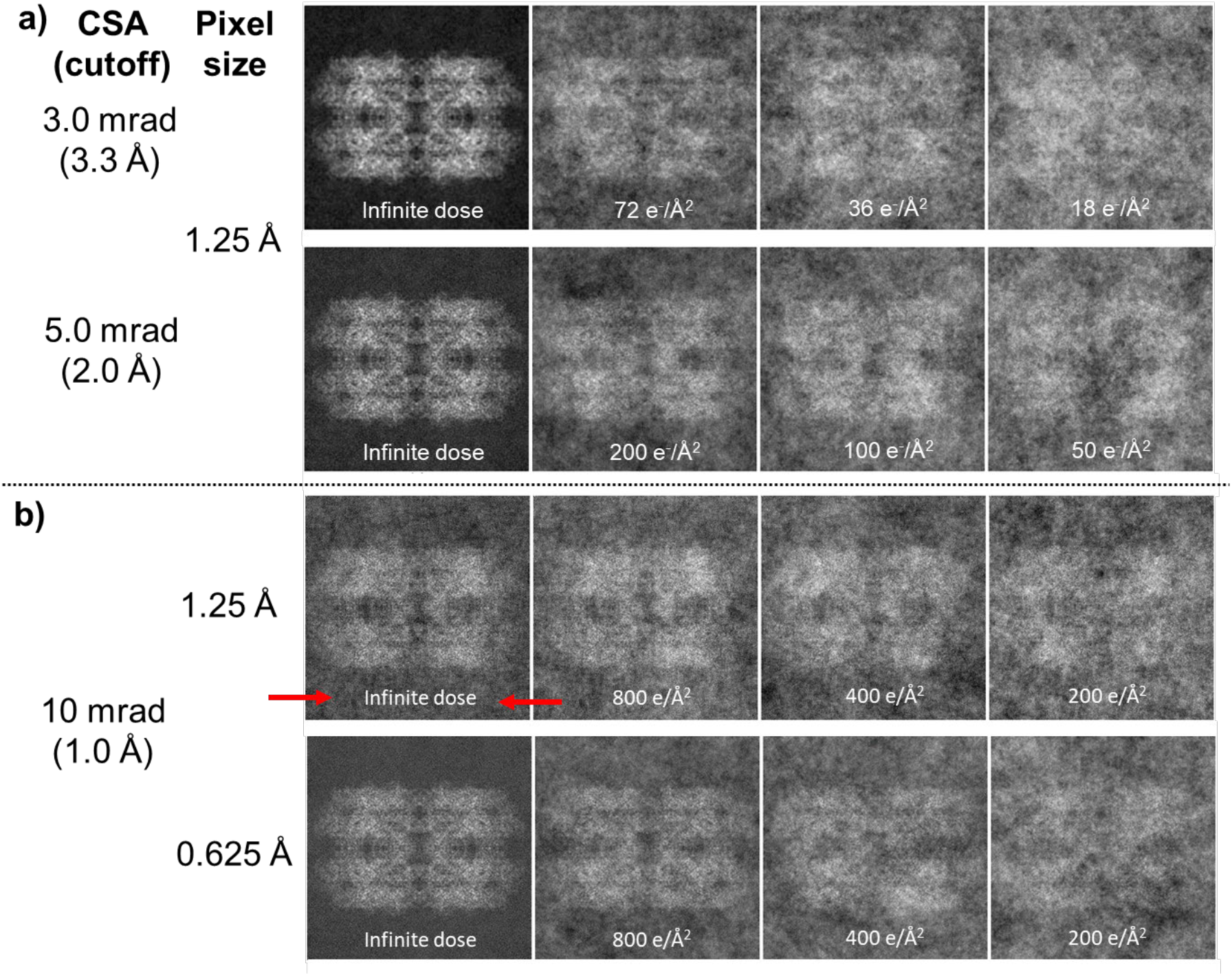
Simulation of iDPC-STEM micrographs of a hemoglobin particle in amorphous ice to illustrate additional limiting effects of larger CSA beam sizes. Imaging conditions used for simulations: voltage 300 kV, image size of 256 × 256 in (**a**) and (**b**) top row, giving pixel size of 1.25 Å, image size of 512 × 512 in (**b**) bottom row, giving pixel size of 0.625 Å. The CSA’s of the beams with corresponding cutoff frequency resolutions and pixel size are indicated on the left. (**a**) Upon increase in CSA at a constant pixel size, the SNR deteriorates due to larger solid angle is covered using the same number of electrons. To maintain the SNRs, higher electron doses can be employed (follow top to bottom). (**b**) Upon increase in CSA at a constant pixel size, the cutoff frequency resolution becomes smaller than the pixel size (here 1.0 Å < 1.25Å at 10 mrad CSA is shown) causing aliasing when frequencies higher than the cut-off frequency fold back into the low-frequency signal (top). The effect of aliasing is most clearly visible at infinite dose (indicated by red arrows). The effect is not present when pixel size smaller than the cut-off resolution is used (bottom).

